# An evolutionary hourglass of herbivore-induced transcriptomic responses in *Nicotiana attenuata*

**DOI:** 10.1101/034603

**Authors:** Matthew Durrant, Justin Boyer, Ian T. Baldwin, Shuqing Xu

## Abstract

Herbivore induced defences are robust, evolve rapidly and activated in plants when specific elicitors, frequently found in the herbivores’ oral secretions (OS) are introduced into wounds during attack. How these complex induced defences evolve remains unclear. Here, we show that herbivore-induced transcriptomic responses in a wild tobacco, Nicotiana attenuata, display an evolutionary hourglass: the pattern that characterises the transcriptomic evolution of embryogenesis in animals, plants, and fungi. While relatively young and rapidly evolving genes involved in signal perception and processing to regulate defence metabolite biosynthesis are recruited both early (1 h) and late (9-21 h) in the defence elicitation process, a group of highly conserved and older genes involved in transcriptomic regulation are activated in the middle stage (5 h). The appearance of the evolutionary hourglass architecture in both developmental and defence elicitation processes may reflect the importance of robustness and evolvability in the signalling of these important biological processes.

## Introduction

Herbivore-induced defences are widespread in plants and play an important role in maintaining plant fitness when they are under attack ^1^. The molecular mechanisms and ecological functions of herbivore-induced signalling cascades and defence responses have been examined in several plant systems ^12^. After attack, chemical cues (herbivore-associated elicitors: HAE) in insect oral secretions (OS) elicit a series of signalling cascades and induced defences in plants, which directly or indirectly deter feeding herbivores and protect plants from further damage^13^. For example, in a wild tobacco, *Nicotiana attenuata,* which is an ecological model plant for studying herbivore-induced defences, the fatty acid conjugates (FAC) found in the oral secretion (OS_*Ms*_) of a Solanaceae specialists herbivore, *Manduca sexta,* elicit rapid phytohormonal changes in leaves, including the rapid accumulation of jasmonic acid (JA) and its derivatives when OS_*Ms*_ are introduced into wounds as larvae feed on leaves^2^,^4^. The amplification of the wound-induced JA burst by OS_*Ms*_ activates the biosynthesis of several potent herbivore toxins, such as phenolamides, 17-hydroxygeranyllinalool diterpene glycosides (HGL-DTG) etc^5,6^, which function as anti-herbivore defences in both the laboratory and the native environment of *N. attenuata*^6-8^.

However, induced defences can also have negative effects on plant fitness due to their physiological and ecological costs^9-12^. In *N. attenuata,* activating induced defences by increasing endogenous JA levels reduces plant fitness by 26% when plants are protected from herbivores^9^. Therefore, the defences in different plant populations and species that grow in heterogeneous environments are subject to divergent selections, which are determined by the effectiveness of induced defences against the local herbivore communities and the costs involved in the activating the induced defences. Indeed, comparative studies on induced defences within and among species have suggested that herbivore-induced defences in plants are highly specific and evolving rapidly^1,13-19^ one plant can elicit different induced defences to different herbivores and different plant species show diverse induced defences to the attack of the same herbivore. Their rapid turnover and common occurance among plant families further suggests that inducing defences and producing them on-demand is a robust strategy for balancing these highly context-specific costs and benefits. However, how herbivore-induced defences evolve, as well as the mechanisms that facilitate their robustness and specificity to diverse and dynamic biotic stresses remains unknown.

Phylotranscriptomic analysis which incorporates the age of the genes into transcriptomic analysis, has proven to be an exceptionally valuble way of understanding the evolution of the developmental processes, displite methodological controversies on gene age estimation^20^. Using this approach, recent studies have shown that transcriptomes of embryogenesis in animals, plants and fungi display an evolutionary hourglass^21-23^, in which genes involved in the early- and late-embryogenic stages are relatively younger and evolve faster than genes from mid-embryogenic stages. One of compelling hypotheses for the establishment and maintenance of such hourglass pattern is their differential interactions with ecological factors at the different developmental stages^22^. For example in animal embryogenesis, while early (zygote) and late (juvenile and adults) embryonic stages often interact with environmental stimuli, the mid-embryonic stages that characterize the phylotypic phase are normally not in direct contact with environment and thus less likely to be subject to ecological adaptations and evolutionary changes^22^. The same logic would predict a similar hourglass pattern in herbivore-induced responses, as only signal perception and the resulting responses are directly in contact to the environment. To test this inference, we analysed the evolution of herbivore-induced transcriptomic responses in *N. attenuata* using phylotranscriptomic approach. The results show that induced-defence signalling indeed displays an evolutionary hourglass. We hypothesize that the hourglass, which reflects modulation and signalling architecture of induced defences may facilitate their evolvability and robustness.

## Results and Discussions

HAE are known to elicit different transcriptomic and metabolomic responses at different times after elicitation in *N. attenuata*^5^, indicating the modulation of HAE-induced defence responses. Such modulations allow us to study the evolutionary patterns of defence signalling with an approach similar to the one used to study the evolution of embryogenesis in animals ‘ and plants by combining transcriptomic induction of genome-wide microarray data with two different evolutionary distance measurements: evolutionary age and sequence divergence. The evolutionary age of each *N. attenuata* gene was estimated with a phylostratigraphic map, constructed by identifying the most distant phylogenetic node that contains at least one species with a detectable homolog^23,24^ In total, 35,096 *N. attenuata* genes were assigned to 13 phylostratigraphic groups (phylostrata; Figure S1), with oldest genes assigned to phylostratum 1 (PS1, shares homologues with prokaryotes) and the youngest genes assigned to PS13 (*N. attenuata* specific). To estimate the sequence divergence, we calculated the Ka/Ks ratio, an indicator of selection pressure at protein coding region, between *N. attenuata* and *N. obtusifolia,* which diverged ∼7 million years ago (MYA)^25^, and between *N. attenuata* and tomato, which diverged ∼ 24 MYA^25^.

To capture the evolutionary properties of genes induced by HAE, we computed two different transcriptomic induction indices for each gene: transcriptomic induction age (TIA), which combines gene induction (log_2_ fold change) and gene age (see Materials and Methods); and transcriptomic induction divergence (TID), which combines gene induction (log_2_ fold change) and sequence divergence. Here, the TIA represents the mean evolutionary age of HAE-induced genes (phylostratum) weighted by its induction level. Similarly, TID represents the mean sequence divergence of HAE-induced genes, where a gene’s sequence divergence (Ka/Ks) is weighted by its induction level (see Materials and Methods). Together, TIA and TID indices provide complementary and independent measurements of evolutionary distances^23^ (Table S2).

TIA and TID were calculated using the microarray data from a HAE-induced 21 h time course experiment sampled at 4 h intervals from locally treated leaves (TL), systemic leaves (SL) and systemic roots (RT), from both control and *M. sexta* oral secretion (*OS_Ms_)* induced wild type *N. attenuata* plants (Figure S2)^26^. The results revealed that HAE-induced transcriptomic responses in *N. attenuata* locally treated leaves display an evolutionary hourglass (Figure 1). At 1 h after elicitation, the induced genes are relatively young (high TIA) and divergent (high TID). At 5 h, the induced genes are dominated by old and conserved genes. And at the later time points (9-21 h), the induced genes are relatively young and highly divergent again. The permutation tests described in Drost et.al.^27^ revealed that the TIA and TID values were significantly different from a flat line (P = 2.95e-54) and consistent with an hourglass pattern (*P* = 1.63e-28), with 1 h as early, 5 h as middle and 9-21 h as late time points (see Materials and Methods). Calculating the TIA and TID for up and down-regulated genes separately reveals that up-regulated genes are primarily responsible for the hourglass (Figure 1). We also found a similar hourglass in the elicitation of systemic leaves (P = 0.025), but not in roots (P = 0.66) (Figure S3). The similar up-regulation of the same genes in systemic and local leaves was responsible for the common hourglass response of these tissues (Table S1).

**Figure 1.**
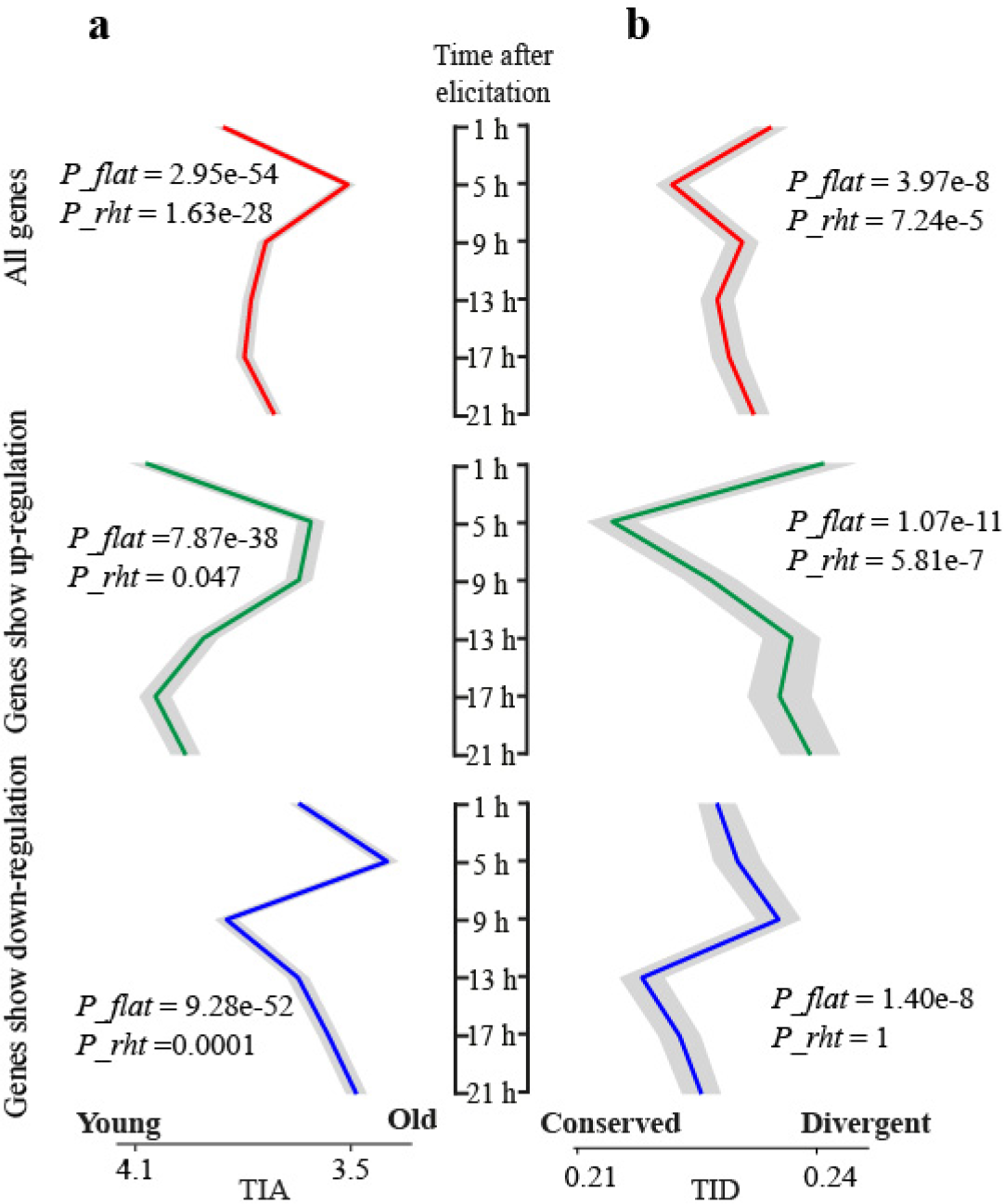
Transcriptome induction age (TIA) and transcriptome induction divergence (TID) in locally treated *N. attenuata* leaves after simulated herbivory. a and b. Red, green and blue lines designate mean indecies calculated on all genes, up- and down-regulated genes, respectively. The grey ribbons refer to standard deviations. P_*_flat*_ and P_*_rht*_ indicate the P value from a flat line and reductive hourglass tests, respectively. P_*_flat*_ less than 0.05 indicates the pattern is significantly different from a flat line and P_*_rht*_ less than 0.05 indicates the pattern follows an hourglass (high-low-high) pattern.

To test the robustness of the observed hourglass pattern, we compared the TIA index based on gene ages estimated using three different homologue searching algorithms, BLASTP, PSI-BLAST and HMMER. We calculated the TID index based on sequence divergences between *N. attenuata* and *N. obtusifolia,* and between *N. attenuata* and *Solanum lycopersicum* (tomato) which diverged ∼ 24 MYA^25^. Our analyses revealed that the observed hourglass pattern was robust to different estimations of gene age and gene divergence (Figure S4 and Figure S5). To capture an additional OS_Ms_-induced early transcriptomic response, we analysed one additional microarray dataset that measured transcriptomic responses at 30 min in TL after OS_Ms_-induction in WT plants that were transformed with an empty vector, since these plants show very similar induced responses and overall phenotypes to WT plants. This analysis revealed that both TIA and TID values were high at 30 min after elicitation and the hourglass pattern was robustly reconfirmed (Figure S6).

We calculated the log odds ratio to identify over-represented phylostratigraphic (PS) groups at different time points for the significantly up-regulated genes (false discovery rate adjusted *P* < 0.05, absolute value of log_2_ fold change > 1) induced by HAE. Consistently, the analysis showed that old genes (PS < 4) were significantly over-represented at 5 h after elicitation (Figure 2). The larger proportion of old genes induced at 5 h was largely due to genes recruited from phylostratigraphic group 2. In addition, genes that were significantly induced by HAE elicitation at 5 h showed lower Ka/Ks ratios between *N. attenuata* and *N. obtusifolia* than genes induced at other time points (Figure S7), consistent with the inference that genes induced at 5 h are more evolutionarily conserved than those genes induced at other time points.

**Figure 2.**
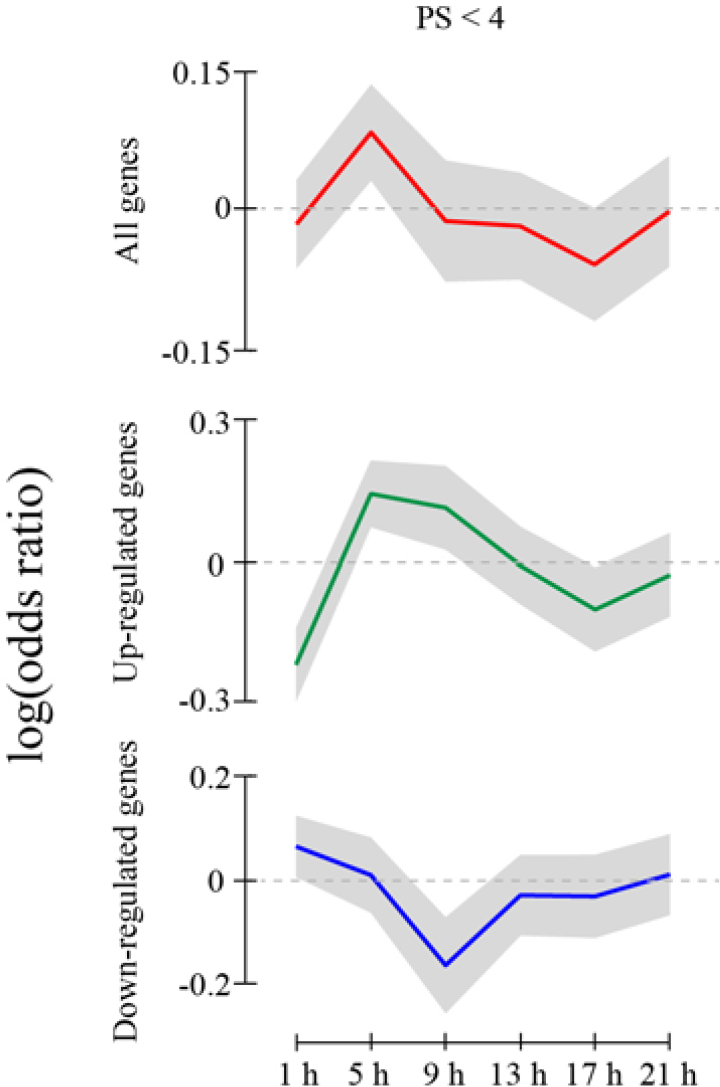
The log odds ratio reveals that old genes are overrepresented 5 h after induction. Red, green and blue lines refer to mean index calculated based on all differentially expressed genes, significantly up-regulated genes, and significantly down-regulated genes with a phylostratigraphic (PS) group less than 4. X-axis indicates the time after induction; Y-axis indicates the log odds ratio. The grey ribbons refer to 95% confidence interval.

To further understand the mechanism underlying the observed evolutionary hourglass pattern, we performed gene ontology (GO) enrichment analysis on the significantly induced genes at each time point (Supplementary dataset 1). At 1 h, the response was enriched in defence and stress signalling processing genes, such as responses to biotic stresses, JA signalling pathways, etc. These genes are known to play key roles in plant-environment interactions and are likely rapidly evolving^28,29^. At 5 h, the genes were enriched in functions related to RNA translation and modification (Figure 3), a highly conserved and central part of the cellular machinery of Eukaryota. Although no specific GO terms were enriched in late-induced genes likely due to their relatively young age and no functional annotations were available, several mediate the biosynthesis of *Nicotiana* specific defence metabolites, such as 17-hydroxygeranyllinalool diterpene glycosides (HGL-DTG), potent herbivore toxins^5, 7^. This evolutionary hourglass thus reflects the architecture and modulation of an HAE-induced signalling cascade in *N. attenuata* leaves (Figure 4).

**Figure 3.**
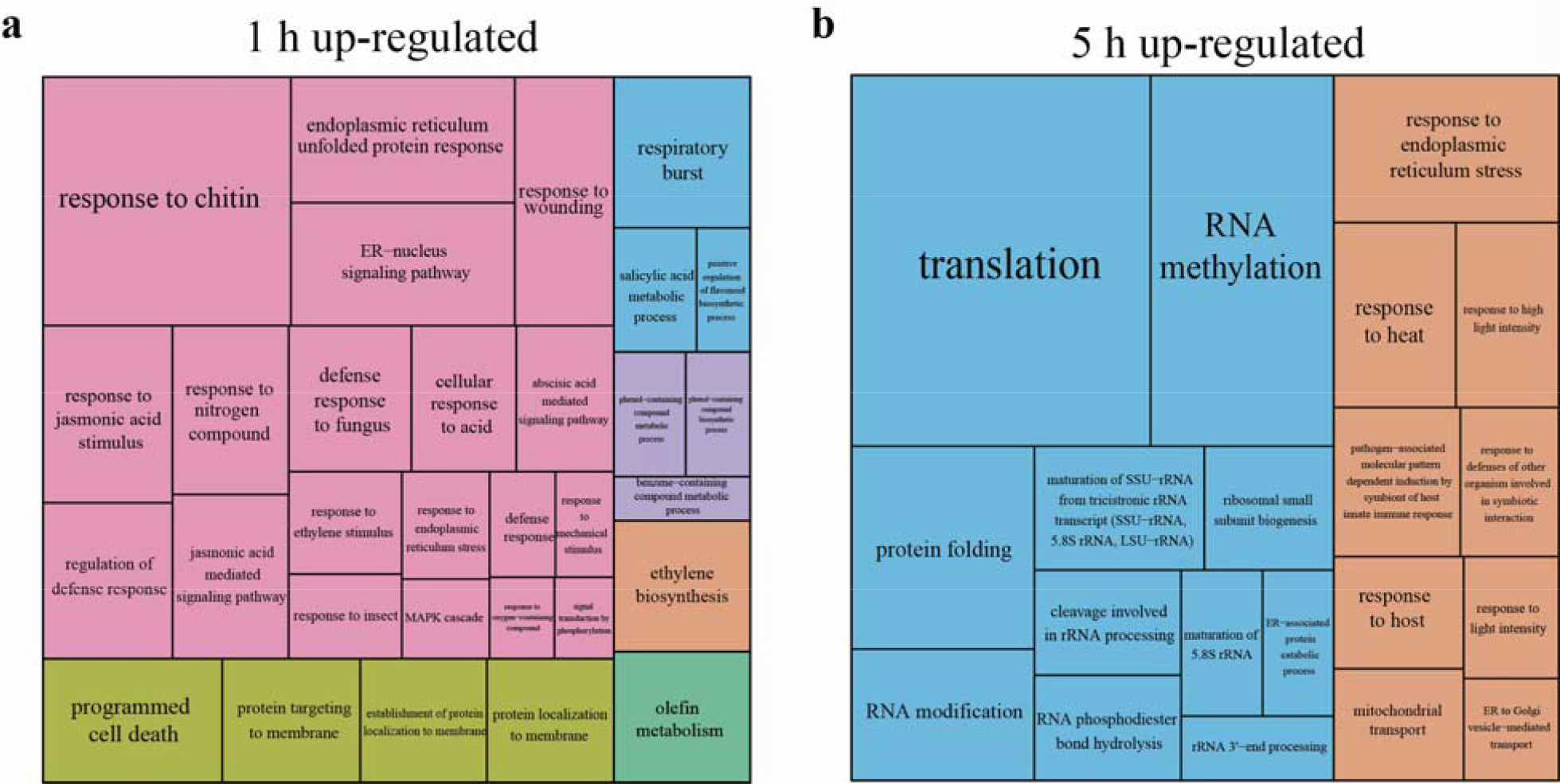
GO terms enriched in genes up-regulated at 1 and 5 h after HAE-induction. While GO terms related to defence signalling are enriched in genes induced at 1 h (left), GO terms related to RNA-modifications and regulations were enriched in genes induced at 5 h (right). The relative size of each GO category is proportional to their statistical significance.

**Figure 4.**
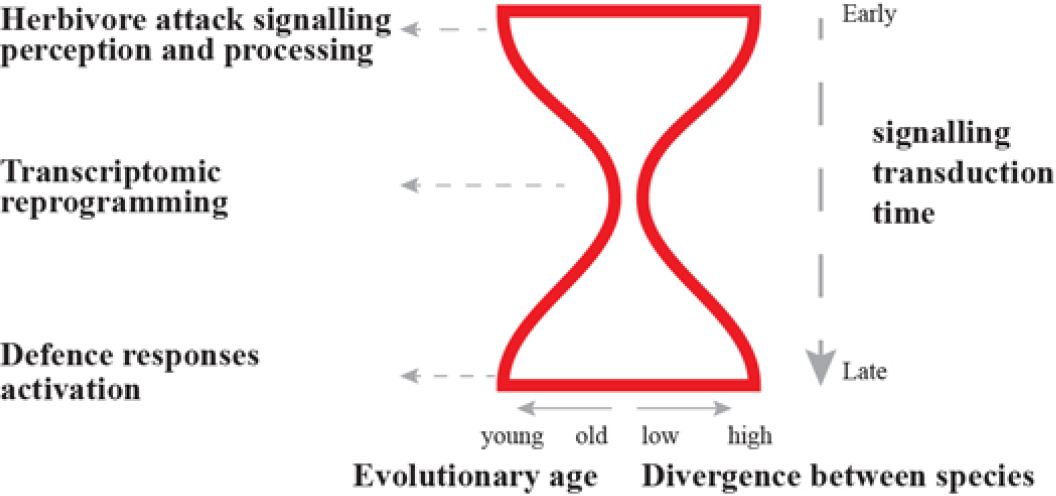
Evolutionary hourglass of herbivore-induced defence signalling in N. attenuata. Herbivore induced defence signalling is modulated in three phases. 1)immediately after a plant perceives herbivore attack, a large group of rapidly evolving and relatively young genes involved in signalling perception and processing are elicited; 2), the perceived and processed signals result in trancriptomic reprograming by activating a group of highly conserved and old genes involved in RNA-regulation machinery; 3) at later time points, the reprogramed transcriptomes activate highly specific defence responses by recruiting relatively young and rapidly evolving genes.

Is the observed hourglass pattern specific to HAE-induced defence signalling? We computed TIA and TID values from locally treated leaves of *Arabidopsis thaliana* induced by flg22, a bacterial associated elicitor^30^. Throughout the 3 h time course of this experiment, both TIA and TID decreased (Figure S8), consistent with the HAE-induced pattern observed at early time points (30 min -5 h) in *N. attenuata.* As data are not available for later times in the elicitation process, we do not know if flg22 elicits the exact same hourglass pattern as HAE elicitation. However, the patterns found in the available data for flg22- and HAE-induced TIA and TID patterns are consistent with a general hourglass response pattem in plant biotic stress-induced defence signalling. Interestingly, the TIA and TID showed different patterns when plants were infected by different living pathogens (Figure S9), likely due to changes in defence signalling caused by pathogen-plant interactions that follow the early pathogen recognition responses.

A strong inference of this analysis is that the different modules in signalling cascades that mediate defence responses respond differently to natural selection. While genes involved in signalling modules that are directly interacting with environmental factors are evolving rapidly, the signalling modules in the middle stage are relatively conserved. Clearly many more resistance responses elicited by different biotic and abiotic stimuli need to be examined with similar phylotranscriptomic approaches in different plant species to test the robustness of the hourglass phenomena. The challenge will be to develop/find/mine datasets that are sufficiently deep in their temporal analysis to capture the complete ontogeny of a discrete response and without having the response be confounded by additional cycles of elicitation and response, as commonly occurs in biotrophic pathogen-plant interactions.

Evolutionary hourglasses describe embryogenesis in animals^22,24^, plants^23^ and fungi^21^. Here, a similar transcriptional hourglass is found in an induced defence response. Although the embryogenesis and induced defences are distinct biological processes, they share the similar feature: the modulation of signalling networks and differential interactions with environmental factors among modules^22-24,31^. Systems biologists have predicted that the modulation and hourglass architecture (bow-tie shape) of signalling networks can facilitate evolvability and robustness of traits^32,33^; both of these features are required for embryogenesis and induced defence. Therefore it is plausible that the modulation of the signalling network itself in induced defences and embryogenesis might be a consequence of adaptations that facilitated their robustness. Synthetic allopolyploid plants may represent an excellent system in which to test this inference. Allopolyploidy, whether it originated in the laboratory or in nature, occurs when the genomes of two different species fuse and new signalling systems are produced that emerge from the recruitment of different modules of the parential species in new combinations^34^. A prediction from this study would be that the variation of induced defences among offspring produced from these interspecies fusions displays a similar hourglass pattern, in which transcriptomic responses at early and late time points after elicitation are more variable than the ones in the middle.

Both developmental processes and herbivore-induced defence responses can also be seen as examples of phenotypic plasticity, in that a single genotype can produce multiple phenotypes in response to signals from the organism’s environment. These phenotypically plastic responses can profoundly influence the process of adaptation, diversification, and more controversially, speciation, as a jack-of-all-trades genotype may impede the speciation/evolutionary process^35^. Understanding the evolutionary history of the different signalling modules that are recruited in these responses may help to resolve some of the controversies, as evolutionarily conserved modules are sandwiched between highly diverging modules in these hourglass patterns. We predict that phylotranscriptomic analyses of developmental signal cascades that mediate phenotypic plasticity would be enriched in hourglass patterns.

## Materials and Methods

### Phylostratigraphic map

To construct the phylostratigraphic map (Figure S1), we used BLASTP from the BLAST (v2.2.25+) suite to search the curated NCBI taxonomy database^22,36^ to assign *N. attenuata* genes to 13 phylostrata. This method is similar to methods used in previous studies^22,36^, with some modifications. In brief, all protein-coding sequences of *N. attenuata* were compared to the non-redundant (nr) NCBI protein database (downloaded on April 29^th^, 2014) by searching BLASTP with an E-value cut-off of^10-3^. The BLASTP results were further filtered to exclude synthetic sequences, viruses, and sequences that do not descend from the ‘cellular organisms’ phylostratum. Due to the scarcity of protein sequences from the Nicotiana genus in the nr database, all *N. attenuata* genes without a match were further searched against a locally stored *N. obtusifolia* genome. A gene was allocated to phylostratum (PS) 12 (Nicotiana specific) if a hit to *N. obtusifolia* was detected or to PS 13 (*N. attenuata* specific) if no hit was detected. All genes were assigned to the phylogenetically most ancient PS containing at least one species with at least one blast hit using a custom python script. This method assumes that genes with shared domains belong to the same gene family, and therefore subsequent duplications of founder genes are generally assigned to the same PS as the founder gene, regardless of the time period in which the duplication event occurred^36^.

Though this method has been used previously^22,36^, a recent study by Moyers and Zhang has demonstrated that using the BLASTP algorithm to find homologs can underestimate a gene’s phylostratigraphic age and result in a biased phylostratigraphic map^20^. To test the robustness of the hourglass pattern, we used two additional homolog search algorithms, PSI-BLAST and PHMMER, which use sequence profile information to search distant homologs. PSI-BLAST (from BLAST 2.2.25+) was run with a cutoff value of^10-3^ for four iterations^37^. HMMER (version: 3.1b1) was run with default parameters and with an E-value cutoff of^10-3^. While the phylostratigraphic map of *N. attenuata* genes based on BLASTP resembles the distribution reported in *A. thaliana* by Quint et al^23^, the *N. attenuata* phylostratigraphic map based on PSI-BLAST resulted in a larger number of genes assigned to earlier PS groups, as predicted by Moyers and Zhang^20^. For example, genes assigned to the PS1 group increased from 4326 (BLASTP) to 6309 (PSI-BLAST). Such shifts in the gene age distribution towards earlier phylostrata on the phylostratigraphic map was even more pronounced when the PHMMER algorithm was used, which resulted in more than a three-fold increase of genes assigned to PS 1 group (from 4326 to 16188 in comparison to BLASTP, Figure S1).

### Ka/Ks ratios

The gKaKs (v1.2.3)^38^ was used to calculate the genome-wide substitution rate between *N. attenuata* and *N. obtusifolia* (diverged ∼ 7 MYA), and between *N. attenuata* and S. lycopersicum (diverged ∼ 24 MYA)^25^. All predicted *N. attenuata* protein coding sequences were used as a query, and the assembled and repeat masked *N. obtusifolia* and S. lycopersicum genomes were used as target genomes. In this pipeline, if the query gene has more than one best match (exact same score) in the target genome, then they were removed to reduce the errors resulted from calculating Ka/Ks from non-orthologous gene pairs. The minimum identity was set to 0.8. The Ka/Ks ratio was calculated using the codeml method from the PAML package (version 4.7). Genes with a Ks value less than 0.05 and greater than 1 were removed from the downstream analysis. The K_a_/K_s_ ratio is an indicator of selection pressure on protein coding genes and thus reflects the natural selection that drives the molecular evolution of analysed genes. Similar to a previous study^23,27^, our analysis showed that the sequence divergence and gene age only show weak correlations (Figure S10) and thus can provide complementary evidence for estimating evolutionary distance.

### Transcriptome induction indices

The transcriptome indices were calculated based on the microarray data^5^ of locally treated leaves, systemic leaves and roots at six time points within 21 h of a simulated herbivore attack (Figure S2). This data set contains two groups of microarray data for each tissue: an herbivore-induced group (wounding + oral secretion (OS) from *M. sexta* to simulate herbivore attack), and a control group (no manipulation). Each group had three biological replicates. The original microarray datasets were obtained from the NCBI gene expression database (BioProject ID: PRJNA143589) and quantile normalization was applied to all microarray data before statistical analysis. The statistical differences and fold change of each gene between control (no manipulation) and the induced group (wounding + oral secretion) for each time point were calculated using the limma (v2.14) package in R (v.3.0.2). For each data point, two different transcriptome indices were calculated: the transcriptome induction age index (TIA), which was calculated based on the expression fold change and gene age; and the transcriptome induction divergence index (TID), which was calculated based on the expression fold change and sequence divergence (K_a_/K_s_). The TIA is a weighted mean age of the transcriptome that is induced at each time point. The TID is a weighted mean evolutionary divergence of the transcriptome that is induced at each time point. The TIA and TID are analogous to the TAI (transcriptome age index) and TDI (transcriptome divergence index) found in previous studies^21,23,24,27^. The TIA and TID are defined as follows:

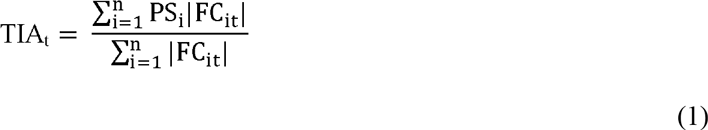

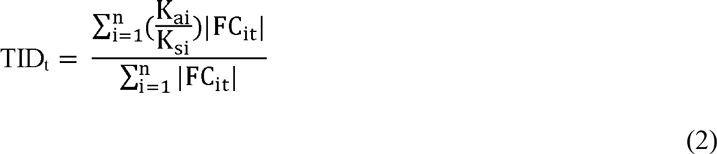

Where t is a time point, n is the total number of genes analysed, PS! is the assigned PS of gene i, FC_lt_ is the log_2_ fold change of gene i at time point t, and 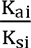 is the 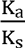 of gene i. In order to compute these two indices, KsiK_s_ all the probes from the microarray were mapped to the predicted protein coding genes using BLASTN, and probes that could be mapped to more than one gene with 60bp matches and 100% identity were removed.

We calculated TIA and TID indices instead of original TAI and TDI indices for two reasons: 1), the main goal of our analysis was to analyse the evolution of induced transcriptomic responses, therefore, the original TAI and TDI that only use gene expression information do not adequately reflect the levels of induction by HAE; 2), the time course experiment lasted 21 h, and the diurnal and circadian rhythms can strongly influence gene expression changes. To minimize the effects of diurnal and circadian rhythms, we calculated induced fold changes based samples collected at same time.

Only genes with at least one unique probe on the microarray dataset were considered for TIA and TID calculations. In total, 17,195 and 12,267 genes were analysed from the microarray data to calculate the TIA and TID, respectively.

For flg22 induced transcriptomic responses in leaves, expression profiles (GSE51720) based on a sequencing technique from a time course experiment that included *Arabidopsis thaliana* leaves induced by flg22 within 3 h were used^30^. The log_2_ fold change data generated from Rallapalli, G. et al^30^, gene age and Ka/Ks information from the development hourglass study by Quint et al^23^ were used for calculating TIA and TID, using similar methods to those mentioned above. For bacterial induced transcriptomic responses, the microarray data from AtGenExpress biotic stress dataset were used (data were originally downloaded from http://www.weigelworld.org/ in October 2013). The log_2_ fold changes were calculated using the R package limma (v2.14). In total, the TIA and TID patterns from induced transcriptome responses induced by four different Pseudomonas strains (Phaseolicola, HrcC, DC300, avrRpm1) and Phytophthora infestans were analysed using the approach described above.

To test whether the TIA and TID values were significantly different from a flat line and consistent with an hourglass pattern, the permutation tests described in Drost et. al^27^ were used (10000 permutations and 100 runs). Because the TIA and TID values differ among later time points, the distance values do not fit a normal distribution, which was assumed to be the case in the orginal method for the calculation of *P* values^27^. Therefore, we used a more conservative approach by calculating *P* values based on only three time points: the time point with the lowest TIA or TID value (for herbivore dataset, 5 h) was assumed to be the middle stage, the lowest TIA or TID value before and after middle time point was selected as early and late stages, respectively. The *P* values calculated using this approach are therefore usually higher (more conservative estimation) than they would be if all time points were included in the later stage sample. The conclusions about the TIA and TID patterns were robust even when using this more conservative approach, and also when usingnon-parametric statistical tests on all different time points.

### Log odds ratio

We calculated the log-odds ratio of significantly induced genes from each phylostrata. The significantly induced genes were identified based on whether their expression was significantly induced by *M. sexta* OS in comparison to control (FDR adjusted P-value less than 0.05 and an absolute log_2_ fold change greater than 1) for each time point. Then the log odds ratio was calculated as:

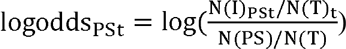

Where PS is phylostrata group, t is each time point; N(I)_PSt_ is the number genes that were induced at the time t; N(T)_t_ is the number of significantly induced genes at time t; N (PS) is total number of genes belong to the phylostrata group PS; N(T) is the informative genes from the microarray. For each PS group, a log-odds ratio greater than 0 indicates that the genes from the tested PS group are over represented among the significantly OS_*Ms*_-induced genes, whereas a log-odds ratio less than 0 indicates under-representation. The confidence intervals for the odds ratio calculated by appealing to the asymptotic normality of the log-odds ratio, which has a limiting variance given by the square root of the sum of the reciprocals of these four numbers. The log-odds ratios were calculated for both merged and separated PS groups. For the merged PS groups, we classified all genes into two groups, young (PS >= 4) and old (PS < 4). To calculate *P* values, a generalized linear model (GLM, with binomial distribution) was used.

### GO enrichment analysis

To further understand the mechanism of the herbivore-induced signalling hourglass pattern, GO enrichment analyses were conducted on the significantly induced genes for each time point. The Cytoscape app ClueGO^39^ was used to determine significant GO groups with an adjusted *P* value less than 0.05. The REViGO online visualization tool^40^ was used to reduce this list of redundant GO categories and to produce tree maps proportioned by the statistical significance of each category.

The original data and the R scripts used for analysing TIA and TID indices are deposited at labarchives (https://goo.gl/DlJ9Zo).

## Acknowledgments

We thank T. Brockmüller and Z. Ling for help with data analysis and discussions. We are grateful for the funding by the Swiss National Science Foundation (project number: PEBZP3-142886 to SX), the Marie Curie Intra-European Fellowship (IEF, project number: 328935 to SX), the Max Planck Society, European Research Council advanced grant ClockworkGreen (project number: 293926 to ITB), and Brigham Young University (travel grants to JB and MD).

